# Effect of concentration and hydraulic reaction time on the removal of pharmaceutical compounds in a membrane bioreactor inoculated with activated sludge

**DOI:** 10.1101/2021.01.29.428761

**Authors:** Ana B. Rios-Miguel, Mike S.M. Jetten, Cornelia U. Welte

**Affiliations:** Department of Microbiology, Radboud University, Institute for Water and Wetland Research, Heyendaalseweg 135, 6525 AJ Nijmegen, The Netherlands; Soehngen Institute of Anaerobic Microbiology, Radboud University, Heyendaalseweg 135, 6525 AJ Nijmegen, The Netherlands

**Keywords:** Organic micropollutants, biodegradation, microorganisms, *Dokdonella*, *Nitrospira*, mobile genetic elements

## Abstract

Pharmaceuticals are often not fully removed in wastewater treatment plants (WWTPs) and are thus being detected at trace levels in water bodies all over the world posing a risk to numerous organisms. These organic micropollutants (OMPs) reach WWTPs at concentrations sometimes too low to serve as growth substrate for microorganisms, thus co-metabolism is thought to be the main conversion mechanism. In this study, the microbial removal of six pharmaceuticals was investigated in a membrane bioreactor at increasing concentrations (4-800 nM) of the compounds and using three different hydraulic retention times (HRT; 1, 3.5, 5 days). The bioreactor was inoculated with activated sludge from a Dutch WWTP and fed with ammonium, acetate, and methanol as main growth substrates to stimulate and mimic co-metabolism in a WWTP. Each pharmaceutical compound had a different average removal efficiency: acetaminophen (100%) > fluoxetine (50%) > metoprolol (25%) > diclofenac (20%) > metformin (15%) > carbamazepine (10%). Higher pharmaceutical influent concentrations proportionally increased the removal rate of each compound, but surprisingly not the removal percentage. Furthermore, only metformin removal improved to 80-100% when HRT or biomass concentration was increased in the reactor. Microbial community changes were followed with 16S rRNA gene amplicon sequencing in response to the increment of supplied pharmaceutical concentration: it was found that *Nitrospirae* and *Planctomycetes* 16S rRNA relative gene abundance decreased, whereas *Acidobacteria* and *Bacteroidetes* increased. Remarkably, the *Dokdonella* genus, previously implicated in acetaminophen metabolism, showed a 30-fold increase in abundance at the highest (800 nM) concentration of pharmaceuticals applied. Taken together, these results suggest that the incomplete removal of most pharmaceutical compounds in WWTPs is neither dependent on concentration nor HRT. Accordingly, we propose a chemical equilibrium or a growth substrate limitation as the responsible mechanisms of the incomplete removal. Finally, *Dokdonella* could be the main acetaminophen degrader under activated sludge conditions, and non-antimicrobial pharmaceuticals might still be toxic to relevant WWTP bacteria.

## 1. Introduction

In the past years, sensitive mass spectrometry methods have enabled the detection of organic pollutants at very low concentrations in nearly all water bodies globally (Jameel et al., 2020). These organic micropollutants (OMPs) include diverse chemicals such as pharmaceutical compounds, and they have detrimental effects on the development, behavior and stress response of aquatic organisms (Gauthier and Vijayan, 2020). Pharmaceutical compounds are emitted into waste streams from different sources (i.e. hospitals, households, industries) and are transported via the sewer system to wastewater treatment plants (WWTPs) where they arrive at very low concentrations compared to more common pollutants such as ammonium and easily degradable organic matter. During the treatment, many of these pharmaceutical compounds are not eliminated from the water and end up in the environment, contaminating surface and ground waters that may ultimately be used as drinking water sources for many organisms, including humans.

Most pharmaceuticals are hydrophilic compounds with low volatility, and are removed to various degrees *via* microbial degradation. Other pharmaceuticals, such as fluoxetine, can also adsorb significantly to biomass (Fernandez-Fontaina et al., 2012; Salgado et al., 2012). Full biodegradation has only been reported in WWTPs for a few pharmaceuticals such as ibuprofen and acetaminophen and several microorganisms have been implicated in their conversion (Żur et al., 2018). On the other hand, many other pharmaceuticals are only partially or not removed at all in WWTPs (i.e. diclofenac, metoprolol). The removal differences between compounds is partially governed by the chemical structure’s properties (Nolte et al., 2020; Nolte et al., 2018). For example, charged compounds are less biodegradable than neutral compounds probably due to a cellular uptake limitation (Nolte et al., 2020; Nolte et al., 2018). In addition to chemical properties, the higher the concentration of pharmaceuticals and other OMPs in the influent of WWTPs, the more these compounds seem to be removed (Nolte et al., 2018; Wang et al., 2020). The same compound can also have different removal rates/efficiencies in different WWTPs which may depend on the operational parameters and the active microbial community present (Kassotaki et al., 2018; Lautz et al., 2017; Wang et al., 2020). For example, nitrifying activities have been correlated to the biodegradation of several OMPs which might be due to the promiscuity of the ammonia monooxygenase enzyme (Fernandez-Fontaina et al., 2012; Men et al., 2017; Tran et al., 2009)

Currently, WWTPs are not very effective at removing OMPs because microorganisms do not share an evolutionary history with these compounds that have only been introduced into the environment in the last decades. Furthermore, evolution seems to be slow due to low concentrations and complexity in chemical structures. Recently, two studies described a putative evolution of acesulfame degradation in WWTPs all around the world (Kahl et al., 2018; Kleinsteuber et al., 2019). Acesulfame was considered recalcitrant in WWTPs until 2014, when several studies started to observe good biodegradation of this compound. Since then more reports showed the decent removal of this compound in WWTPs. In follow-up studies, microorganisms able to use acesulfame as carbon source were isolated (Kahl et al., 2018; Kleinsteuber et al., 2019). This example demonstrates the on-going adaptation of microorganisms towards OMP biodegradation in WWTPs. Mobile genetic elements (MGEs) have been previously suggested to be involved in microbial adaptation. MGEs may thus have played a role in the biodegradation of xenobiotics or organic micropollutants in different environments such as soils and WWTPs (Rios-Miguel et al., 2020; Top and Springael, 2003). Furthermore, previous studies found a correlation between pesticide exposure and MGE abundance in soils and agricultural WWTPs (Dealtry et al., 2014; Dunon et al., 2013). Consequently, MGE abundance might also be correlated to the pharmaceutical concentration entering WWTPs and the subsequent adaptation to degrade OMPs.

Membrane bioreactors (MBRs) are known to perform well in the removal of pharmaceutical compounds (Goswami et al., 2018; Prasertkulsak et al., 2016), sometimes even better than conventional activated sludge systems (Reif et al., 2011; Verlicchi et al., 2012). MBRs are able to separate the solid retention time (SRT) and the hydraulic retention time (HRT), thus keeping slow growing bacteria at high SRT while treating wastewater at regular HRT. In general, higher SRTs (with a maximum of 10-15 days) have been associated to better removal of pharmaceutical compounds (Achermann et al., 2018; Clara et al., 2005; Joss et al., 2005; Nguyen et al., 2020). For some compounds, however, extremely high SRTs are needed such as for diclofenac (more than 100 days) and metoprolol (60 days) (Clara et al., 2005; Fernandez-Fontaina et al., 2012; Gurung et al., 2019). Previous studies also observed a positive correlation between HRT and the removal of some pharmaceuticals (i.e. fluoxetine) (Fernandez-Fontaina et al., 2012). However, recent experiments and models suggested that pharmaceutical and other OMP removal might not improve beyond a specific HRT (Boonnorat et al., 2019; Gonzalez-Gil et al., 2018; Jiang et al., 2018). Furthermore, Gonzalez-Gil et al. hypothesized that this might be due to an equilibrium in the transformation or sorption processes (Gonzalez-Gil et al., 2019; Gonzalez-Gil et al., 2018)

In order to get more insights about the (in)complete degradation of some pharmaceuticals in WWTPs, we have studied the effect of concentration and HRT on pharmaceutical removal in a membrane bioreactor. We also determined the changes in the microbial community in response to higher pharmaceutical concentrations and finally, we tested whether higher pharmaceutical concentrations resulted in higher relative abundance of specific MGEs previously correlated to pesticide occurrence and degradation. In our experimental setup, we inoculated a membrane bioreactor with activated sludge from a Dutch WWTP and fed it with ammonium, acetate, and methanol as main growth substrates. We kept all parameters constant except for the influent pharmaceutical concentration (4-800 nM) or HRT (1, 3.5, 5 days). We monitored pharmaceuticals by liquid chromatography–mass spectrometry (LC-MS), microbial community changes by bacterial 16S rRNA gene amplicon sequencing and MGEs by qPCR.

## 2. Methods

### 2.1. Pharmaceutical selection and bioreactor operation

Six pharmaceuticals were selected based on (i) presence in priority lists, (ii) variability in primary biodegradation rate and (iii) non-antimicrobial characteristics to avoid community changes based on previously known toxicity. The selected pharmaceuticals were acetaminophen, diclofenac, metoprolol, carbamazepine, metformin, and fluoxetine. They were all purchased in solid phase form Merck (Darmstadt, Germany), dissolved in methanol and diluted in water to obtain a 1:1 methanol water solution.

A 5L membrane bioreactor (Applikon Biotechnology B.V., Delft, The Netherlands) was inoculated with 100-times diluted activated sludge from a municipal WWTP in Groesbeek, the Netherlands. The bioreactor was fed with synthetic wastewater containing ammonium (~3 mM), acetate (~3 mM) and methanol (~1 mM) as main energy sources. The detailed composition of the medium can be found in the supplement SI1. The medium was autoclaved and then, pharmaceuticals were added. After an acclimatization period of 180 days at a hydraulic retention time (HRT) of 3.5 days and a solid retention time (SRT) of 15 days, the pharmaceutical concentration in the influent was increased every two weeks from 4 nM to 800 nM. Methanol concentration in the influent did not change, as stock solutions with increasing pharmaceutical concentrations were used. During the process, pharmaceuticals were monitored by LC-MS/MS and the microbial community was analysed by 16S rRNA gene amplicon sequencing. Furthermore, the experiment was run in the dark at constant mechanical stirring (200 rpm), pH = 7, dO_2_ = 1 mg/L, room temperature (20 +/-1 °C) and total suspended solids (TSS) = 0.2-0.3 g/L. Dissolved oxygen (dO_2_) and pH sensor probes were connected to an ADI 1010 controller (Applikon Biotechnology B.V., Delft, The Netherlands) that activated a KHCO_3_ base pump to control the pH and regulated the O_2_/CO_2_ entrance flow in the bioreactor. Nitrogen (ammonium, nitrite, and nitrate) and carbon (acetate and methanol) balances were measured and calculated during the experiment. After this first experiment, pharmaceutical removal was measured at HRTs of 5 and 1 days keeping pharmaceutical concentration at 800 nM.

### 2.2. Carbon, nitrogen, and total suspended solids measurements

Ammonium, nitrite and nitrate assays were performed with technical duplicates in 96-well microtiter plates using a plate reader as previously described in (van Bergen et al., 2020). Briefly, ammonium was measured colorimetrically at 420 nm after reaction with OPA reagent (0.54% (w/v) ortho-phthaldialdehyde [OPA], 0.05% (v/v) β-mercaptoethanol and 10% (v/v) ethanol in 400 mM potassium phosphate buffer (pH 7.3)). Nitrite was measured at 540 nm using the Griess assay. Afterwards, an incubation with vanadium chloride at 60°C reduced all nitrate to nitrite and the sample was measured again at 540 nm. Methanol was measured colorimetrically with the 2,2’-azino-bis-(3-ethylbenzothiazoline-6-sulfonic acid) (ABTS) assay explained elsewhere ((Mangos and Haas, 1996) and https://www.sigmaaldrich.com/technical-documents/protocols/biology/enzymatic-assay-of-alcohol-oxidase.html). We adjusted the volumes to fit in a 96-well plate. First, a stock solution was prepared with the following composition: 14 mL ABTS solution, 100 μL of peroxidase solution (1 mg/mL), 3 mL of 100 mM potassium phosphate buffer pH 7.5, and 100 μL of alcohol oxidase solution (1 mg/mL). Afterwards, 170 μL of the stock solution and 20 μL of samples or standard were added to each well. The plate was incubated for one hour in the dark at room temperature and measured at 405 nm in the plate reader. Finally, acetate was measured by GC-MS using the protocol described in a previous study (in ‘t Zandt et al., 2018). Briefly, 200 μL of 100 mM pentafluorobenzylbromide solution in acetone, 40 μL of 0.5 M phosphate buffer pH=6.8, and 40 μL of sample/standard were mixed in an Eppendorf tube and incubated for one hour at 60 °C. Afterwards, 400 μL of 0.1 mM methylstearate solution in hexane were added and the mixture was vortexed and centrifuged for one minute at maximum speed. The top layer was transferred to a GC vial and measured in a GC-MS.

TSS and VSS were measured as in (van Bergen et al., 2020). Briefly, 40 mL of sample was passed through a 0.45 μm pore size glass-fiber filter, dried overnight in the oven at 105 °C, and then 4 hours at 550 °C. Filters were weighed in the beginning and after each step (105 and 550 degrees) to calculate the number of TSS and VSS, respectively.

### 2.3. Pharmaceutical measurements via LC-MS/MS and mass balances

Samples (2 mL) were taken every week from the medium (influent) and the bioreactor (effluent) in technical triplicates. Samples were centrifuged at maximum speed and the supernatant was immediately treated and analysed as in (van Bergen et al., 2020). Briefly, 0,5 mL of sample was used for liquid-liquid extraction of metformin. One mL of sample was used for solid phase extraction (Oasis HLB 3cc SPE cartridges, sorbent bed of 60 mg from Waters Corporation, Milford, USA) of the other five pharmaceuticals. The extracts were directly injected to an LC-MS system together with the calibration standards prepared in the same way as samples. Internal standards (deuterated compounds) were added to the samples and standards before the extractions. Only parent compounds were determined.

The removal efficiency or percentage of a compound was calculated using equation 1. C_i_ corresponds to the pharmaceutical concentration in the influent and C_e_ in the effluent of the bioreactor (nM).

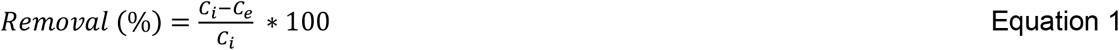

The removal rate of a compound was calculated using equation 2, where TSS corresponds to the total suspended solids (g/L) inside the bioreactor. In our experiment, TSS corresponds to the biomass content.

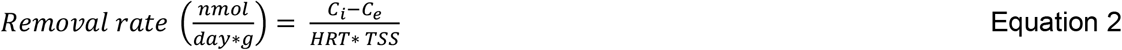

The sorption of fluoxetine was calculated using equation 3 (Ternes et al., 2004), where K_d_ represents the solid-water distribution coefficient and SS the quantity of sludge generated per unit of wastewater treated. The K_d_ value used in this study was taken from (Fernandez-Fontaina et al., 2012).

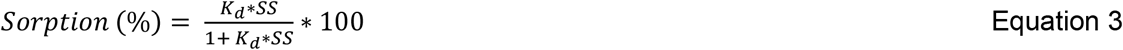

### 2.4. Molecular analysis

#### 2.4.1. Sampling and DNA extraction

Samples (8 mL) were taken every week from the bioreactor in technical triplicates. The samples were centrifuged at maximum speed and the pellet was stored at −20 °C until DNA extraction. DNA was extracted and purified using the DNeasy PowerSoil kit (QIAGEN Benelux B.V.) following the manufacturer’s instructions. The DNA concentration was determined using the Qubit dsDNA HS Assay kit (Thermo Fischer Scientific, Waltham, MA USA) and a Qubit fluorometer (Thermo Fischer Scientific, Waltham, MA USA).

#### 2.4.2. Bacterial 16S rRNA gene sequencing and data analysis

DNA samples were submitted to Macrogen (Seoul, South Korea) for amplicon sequencing of the bacterial 16S V3 and V4 region using Illumina Miseq Next Generation Sequencing. The primers used for amplification were Bac341F (5′-CCTACGGGNGGCWGCAG-3′) and Bac785R (5′-GACTACHVGGGTATCTAATCC-3′) (Klindworth et al., 2013). Subsequent analysis on the sequencing output files was performed within R version 3.4.1 (Team, 2012). Pre-processing of the sequencing data was done using the DADA2 pipeline (Callahan et al., 2016). Taxonomic assignment of the reads were up to the species level when possible using the Silva non-redundant database version 128 (Yilmaz et al., 2014). Count data was normalized to relative abundances. Data visualization and analysis were performed using phyloseq and ggplot packages (McMurdie and Holmes, 2013; Wickham and Wickham, 2007). Chao1, Simpson and Shannon diversity indexes were calculated using the estimate richness function of the phyloseq package. All sequencing data were submitted to the GenBank databases under the BioProject ID PRJNA670630 (reviewer link: https://dataview.ncbi.nlm.nih.gov/object/PRJNA670630?reviewer=dt2eeti6d8l048a7r6t9efict4).

#### 2.4.3. Comammox *amoA* gene PCR

The primers and PCR conditions used to amplify the *amoA* gene of comammox bacteria are explained in Pjevac et al. (Pjevac et al., 2017). Two degenerate primer pairs were used to target the *amoA* gene from clade A and clade B comammox (Table S2). The oligonucleotide primers were obtained from Biolegio (Nijmegen, The Netherlands). PCR reactions were performed in a total volume of 25 μL which contained 1 μL DNA sample of 1 to 10 ng/μL, 0.5 μL of each degenerate primer solution of 20 μM, 12.5 μL of PerfeCTA Quanta master mix (Quanta Bio, Beverly, MA) and 10.5 μL of autoclaved MiliQ water. The PCR protocol consisted of an initial denaturation step at 94°C for 5 min, followed by 25 cycles of denaturation at 94°C for 30 s, primer annealing at 52°C for 45 s, and elongation at 72°C for 1 min. Finally, the last step was an elongation at 72°C for 10 min.

#### 2.4.4. Quantitative PCR

16S rRNA genes were amplified using the above primers. Amplification of the *trfA* gene from IncP-1 α, β, ε, γ, and δ plasmids and the *tnpA* gene from the IS*1071* insertion sequence was performed with primers previously used in (Bahl et al., 2009; Dunon et al., 2013; Providenti et al., 2006). A summary of all primers used in this study can be found in Table S2. All qPCR reactions were performed using 96-well optical PCR plates (Bio-Rad Laboratories Veenendaal, the Netherlands) with optical adhesive covers (Applied Biosystems, Foster City, CA) in a C1000 Touch thermal cycler equipped with a CFX96 Touch™ Real-Time PCR Detection System (Bio-Rad Laboratories, Veenendaal, the Netherlands). The qPCR total volume was 25 μL containing 1 μL DNA sample of 1 to 10 ng/μL, 0.5 μL of each primer solution of 20 μM, 12.5 μL of PerfeCTA Quanta master mix (Quanta Bio, Beverly, MA) and 10.5 μL of autoclaved MiliQ water. Negative controls were added to each run by replacing the template with sterile Milli-Q water. All qPCR data was analyzed using the Bio-Rad CFX Manager version 3.0 (Bio-Rad Laboratories, Veenendaal, the Netherlands). The qPCR protocol consisted of initial denaturation at 95°C 3 min followed by 39 cycles of denaturation, annealing, and extending (95°C 30 sec, 60°C 30 sec, and 72°C 30 sec, respectively), and two final steps at 65°C 5 sec and 95°C 50 sec.

## 3. Results and discussion

### 3.1. Bioreactor performance

The bioreactor performance was monitored throughout the whole experiment using a complementary array of methods (Table S3). Total suspended solids (TSS) were measured every week (Figure S1) and were stable at 0.2-0.3 g/L after the start-up period. Only when the HRT was reduced to 1 day, TSS increased to 0.9 g/L. Activated sludge was diluted around 100x times when inoculated to the bioreactor so that volatile suspended solids (VSS) were similar to TSS (data not shown). Ammonium, acetate and methanol were fully consumed at all measured times (Figure S2). Nitrate was measured in the effluent at concentrations between 500 and 800 μM, indicating that ammonium was partly converted to nitrate during the experiment. The remaining ammonium was likely assimilated. When HRT was reduced to 1 day, nitrite slightly accumulated (~50 μM) and nitrate concentration considerably increased in the effluent (~1.6 mM). This could indicate an imbalance between ammonia- and nitrite-oxidizers and a decreased use of nitrate for assimilation. Pharmaceuticals, 16S rRNA gene amplicons, and mobile genetic element (MGE) monitoring are addressed in the next sections.

### 3.2. Pharmaceutical removal at increasing concentrations

After the start-up period, the bioreactor was close to a steady-state where all variables remained stable throughout the operating period: temperature, dO2, pH, TSS, HRT, SRT, and influent concentrations. When the steady state was reached, the concentration of the pharmaceutical compounds was step-wisely increased in the influent every two weeks, from 4 nM to 800 nM. Removal percentages of pharmaceuticals under different concentration regimes are shown in Figure 1. Overall, each pharmaceutical had a different removal percentage. Acetaminophen was the only compound that was fully removed (100%). Metoprolol and diclofenac removal was between 20% and 40%, carbamazepine removal values varied between 5% and 20%, and metformin removal was between 5 % and 30%. Fluoxetine removal was highly variable, between 25% and 70%, and the influent concentration was around 10 times lower than expected, probably due to its high sorption behaviour to several surfaces, including glass (Peake et al., 2015). For pharmaceuticals with K_d_ (solid-water distribution coefficient) values lower than 0.5 L/g, sorption is considered insignificant in WWTPs (Ternes et al., 2004; Wang et al., 2020). For this reason, sorption was only considered as a removal mechanism for fluoxetine and it was calculated using Equation 3. The K_d_ coefficient value (~1.60 L/g) was taken from a previous study where adsorbed fluoxetine levels were directly measured in nitrifying sludge (Fernandez-Fontaina et al., 2012). An average value of 0.21 g/L of TSS was maintained in the bioreactor at a SRT = 15 days and HRT = 3.5 days. Thus, the amount of biomass generated per litre of medium treated was 0.049 g/L, which corresponds to 7% of fluoxetine sorption. Volatilization was not considered as a removal mechanism based on the low values of Henry’s law constants of the selected pharmaceuticals (taken from the Hazardous Substances Data Bank) and based on previous studies (Suárez et al., 2008).

**Figure 1.**
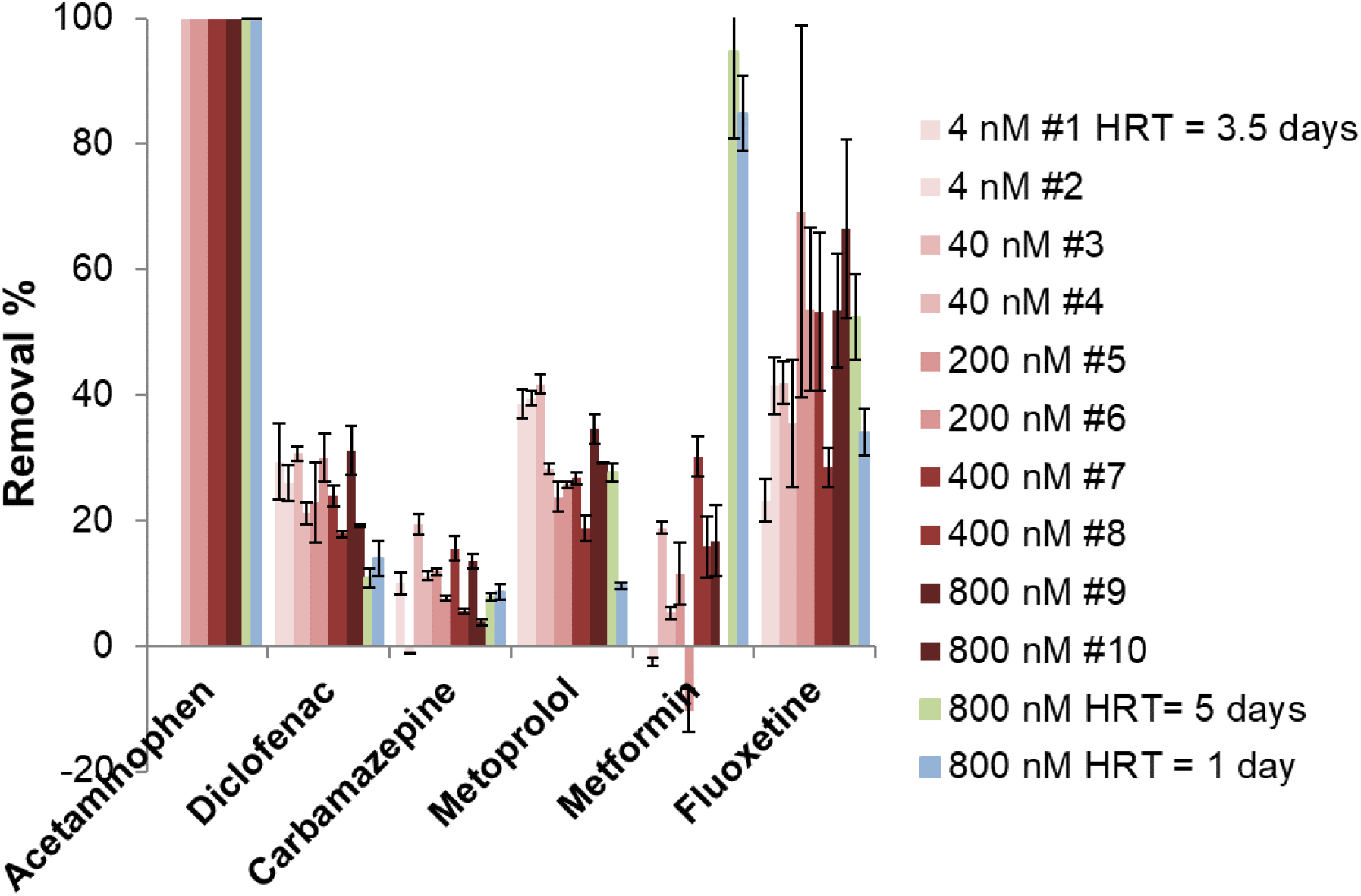
Pharmaceutical removal percentage at different influent concentrations and hydraulic retention times (HRT). Fluoxetine concentration was measured in the influent ten times lower than what it is shown in the graph. Error bars are calculated from technical triplicates. The symbol “#” followed by a number means the week the measurements were done since the starting of the concentration increase experiment.

The removal efficiencies reported in Figure 1 are in line with previous studies, except for metformin, whose removal in WWTPs has been reported much higher (usually more than 80%)(Briones et al., 2016; Oosterhuis et al., 2013). Acetaminophen is known to be removed very well and carbamazepine is known for its high persistence in WWTPs all around the world (Falås et al., 2016; Joss et al., 2005; Wang et al., 2020). Metoprolol and diclofenac removal is variable in WWTPs, but their removal is often slightly higher than carbamazepine removal (Falås et al., 2016; Wang et al., 2020). Diclofenac removal during nitrification has been recently reported to be higher (~84%) than during heterotrophic processes or conventional activated sludge treatment (~50%)(Tran et al., 2009; Wu et al., 2020). However, earlier studies did not observe such high removal of diclofenac in nitrifying sludge, only when solid retention time was extended to 150 days removal was improved to 70% (Fernandez-Fontaina et al., 2016; Fernandez-Fontaina et al., 2012). It might be that rare species present in the sludge proliferated during the high SRT, and then started to contribute to metabolize diclofenac. This might explain the differences in removal under nitrifying conditions. Previous studies showed full removal of metoprolol after four days in activated sludge (Rubirola et al., 2014; van Bergen et al., 2020) and after 10-20 days in constructed wetland biomass (He et al., 2018). Furthermore, the former study also observed an enhancement of metoprolol removal with more dissolved organic matter and a decrease in removal during nitrification. This was further confirmed by a different experiment with partial nitritation-anammox biomass where metoprolol removal was higher under aerobic heterotrophic conditions than under nitrifying conditions (Kassotaki et al., 2018). Furthermore, the addition of allylthiourea (an inhibitor of ammonia monooxygenase and other monooxygenases) did not result in a significant decrease of metoprolol removal. Consequently, metoprolol was probably removed by heterotrophic bacteria in our bioreactor. Furthermore, the removal variability of diclofenac and metoprolol in WWTPs might be due to the differences in nitrification and heterotrophic microbial activities. Fluoxetine has a high K_d_ and its sorption to biomass has a great contribution to the removal percentage in activated sludge (Fernandez-Fontaina et al., 2012). Besides sorption, fluoxetine may also be degraded in nitrifying and heterotrophic cultures (Fernandez-Fontaina et al., 2016; Velázquez and Nacheva, 2017). Therefore, fluoxetine removal in our bioreactor probably happened via sorption and transformation by nitrifiers and heterotrophic bacteria.

The removal percentage of each pharmaceutical did not correlate with the influent concentration (Figure 1). Only small fluctuations in removal percentage were observed for each pharmaceutical. In the case of metoprolol, we observed that higher concentrations slightly decreased the removal percentage. However, when the highest concentration was added to the bioreactor (800 nM), metoprolol removal increased again. In summary, we did not find any effect of concentration on pharmaceutical removal percentage under the conditions tested in our membrane bioreactor. These results are in contradiction to other studies reporting a correlation between influent concentration and removal percentage of some OMPs in WWTPs (Nolte et al., 2018; Onesios-Barry et al., 2014; Wang et al., 2020). Onesios-Barry et al. reported a negative correlation between concentration and removal for pharmaceuticals such as fluorouracil in laboratory columns inoculated with wastewater effluent. The lower removal values at higher concentrations might be related to toxicity. On the other hand, Nolte et al. and Wang et al. reported a positive correlation between OMP concentration and removal. Their data came from several WWTPs with many different variables, whereas our experiment relied on a membrane bioreactor in steady state. High OMP concentrations over long periods of time (years) in WWTPs may lead to microbial adaptation, and consequently, to an increase of removal. In our bioreactor experiment, we increased concentration every two weeks, so we did not expect microbial evolution to occur towards pharmaceutical utilization in this short time frame. Furthermore, the putative toxicity of some pharmaceuticals did not result in lower removal efficiencies probably because the microbial community composition changed during the increasing concentrations (Figure 3) and mitigated this effect.

Our results suggest that the incomplete removal of pharmaceuticals is not dependent on concentration. Consequently, we could rule out hypotheses that rely on threshold concentrations as determinants of partial removal. One example is the cessation of transformation reactions due to product inhibition or toxicity, which would be concentration-dependent. As we did not observe consistent inhibition at one particular pharmaceutical concentration, but rather a similar removal percentage at all tested concentrations, it is unlikely that product inhibition or toxicity was an important factor for the pharmaceuticals in our experiment. The hypothesis of product inhibition for governing the incomplete degradation of OMPs was also discarded in a previous study using a mechanistic modelling approach (Gonzalez-Gil et al., 2018). Another hypothesis for explaining the cessation of transformation reactions relies on catalytic constants of enzyme activity, such as enzyme affinity and corresponding reaction velocity. When a pollutant is transformed, its concentration decreases to a point that might be below the substrate affinity of the responsible enzyme. However, removal percentage would have increased with pharmaceutical concentration if this hypothesis was true. As a result, our experimental evidence does not support these hypotheses where the biodegradation efficiency is dependent on the influent pharmaceutical concentration under concentration regimes relevant to real-world scenario’s.

Although no correlation was found between pharmaceutical influent concentration and removal percentage, the absolute amount of nmols removed increased proportionally with the concentration (Figure 2). This indicates that enzyme saturation was not yet reached and that bacteria had the capacity to transform more pharmaceutical molecules per time even though they did not fully convert them at lower concentrations. Previous studies reported higher biodegradation rates at higher OMP concentrations (in the nM and μM range) in activated sludge, biofilm reactors, pure enzymes, and bacterial isolates (Leng et al., 2016; Lonappan et al., 2017; Svendsen et al., 2020; van Bergen et al., 2020). In many cases, they used the Michaelis-Menten model to describe the increase in biodegradation rates with concentration. This model corresponds to a linear relationship between initial concentration and velocity up until the system saturation, when biodegradation rates do not increase as fast as before and they finally reach a limit. In our bioreactor experiment, the system did not reach saturation probably because pharmaceutical concentrations were increased every two weeks, thus leaving time for the microbial community to shift and adapt.

**Figure 2.**
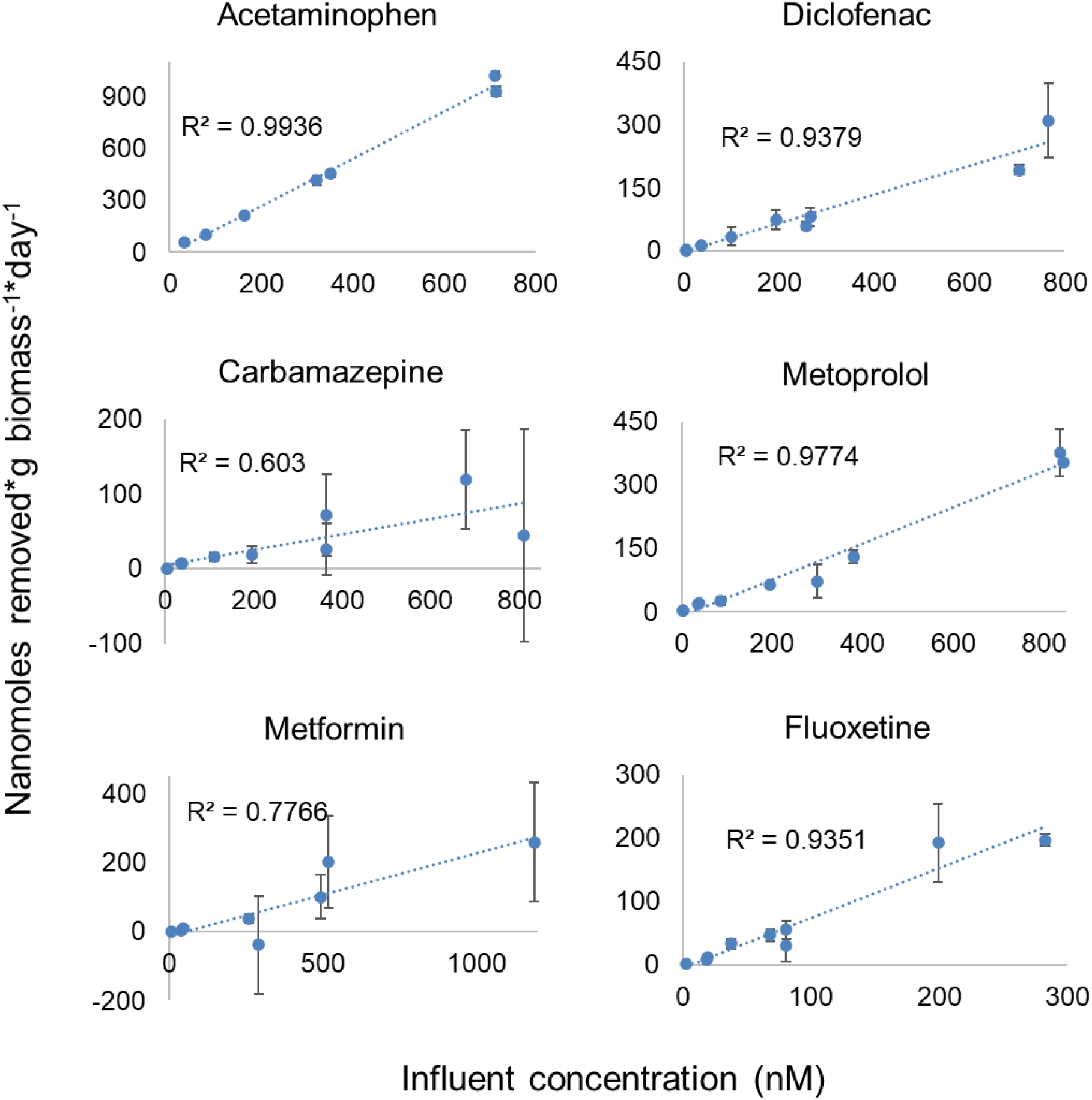
Pharmaceutical removal rate at increasing influent concentrations.

### 3.3. Pharmaceutical removal at different hydraulic retention times

During a second experiment in the same bioreactor, the HRT was increased to five and decreased to one day to prolong/shorten the reaction time of the pharmaceuticals with the microbial community. Influent concentration of pharmaceuticals was kept at 800 nM and operational parameters in the bioreactor remained unchanged. However, biomass concentration increased 3 times to 0.9 g/L at HRT = 1 day due to the increased loading rate of growth substrates, mostly acetate and methanol (Figure S2). Furthermore, higher nitrite and nitrate concentrations were observed at HRT = 1 day (Figure S2). Figure 1 shows the removal percentage of the six pharmaceuticals at different HRT. Acetaminophen was always removed 100% with no effect of HRT on removal percentage, which means that total removal happened in the first 24 hours. HRT had no effect on carbamazepine and fluoxetine removal efficiencies as they were in the same range as with HRT of 3.5 days. Diclofenac removal was slightly lower at HRT 1 and 5 days. Metoprolol removal did not change by increasing HRT to 5 days, but it was lower at HRT = 1 day. Therefore, 10% of the removal happened in the first 24 hours and the remaining 20% in the following 2.5 days reaching a removal maximum of 30%. Metformin was the only pharmaceutical removed to a higher degree by increasing the HRT to 5 days (~95 %) suggesting that the length of the reaction time was responsible for the incomplete removal of metformin in our bioreactor. However, decreasing the HRT to 1 day maintained a high metformin removal percentage of 80% which indicates that another factor was affecting the removal. Biomass concentration at HRT = 1 day was three times higher than in the other HRTs, so this might be the reason why removal was higher. These results suggest that metformin removal depends on the reaction time or HRT and the biomass concentration present in the bioreactor. Metformin is usually very well degraded, so its removal efficiency did not increase with HRT in WWTPs and constructed wetlands (Auvinen et al., 2017; Oosterhuis et al., 2013). Biomass concentration in WWTPs is usually higher than what we used in our experiments, indicating that the data obtained under HRT = 1 day with higher biomass concentrations are more representative of the *in situ* WWTP situation.

Previous studies reported higher removal percentages at higher HRTs for diclofenac and carbamazepine in constructed wetlands (from 2 to 6 days), for fluoxetine in nitrifying and activated sludge (from 0.75 to 1 and 5 days), and for metoprolol in activated sludge (from 1 to 24 hours) (Alvarino et al., 2014; Auvinen et al., 2017; Fernandez-Fontaina et al., 2012; Maurer et al., 2007). However, other studies in activated sludge and anaerobic reactors did not observe an increase in diclofenac removal percentage at higher HRT (at 19-61 hours and 5-11 days, respectively) (Oosterhuis et al., 2013; Queiroz et al., 2012). A recent study in bioaugmented activated sludge showed an increase in diclofenac and carbamazepine removal percentage from 12 to 18 hours, but removal was no longer improved at 24 hours (Boonnorat et al., 2019). These results suggest that increasing the HRT of wastewater bioreactors might increase the removal efficiency of specific pharmaceuticals, but only up to a specific point. This indicates that there will always be a removal constrain not related to kinetics. For example, in our membrane bioreactor this point was 3.5 days for metoprolol and 1 day or lower for carbamazepine. Furthermore, we observed a decrease in diclofenac removal percentage at higher HRT (5 days) which might be related to a desorption or back transformation from diclofenac reaction products.

The incomplete removal of diclofenac, carbamazepine, metoprolol, and fluoxetine was not dependent on concentration nor HRT, so only few removal mechanisms are possible. One hypothesis is that (1) transformation or sorption reactions are reversible and reach an equilibrium. In such a case we would always observe the same removal percentage at different concentrations and no removal increase with time or HRT. This was recently proposed by Gonzalez-Gil et al. using a mechanistic model based on the reversibility of bisphenol phosphorylation by two key enzymes in anaerobic digestion: hexokinase and acetate kinase (Gonzalez-Gil et al., 2019; Gonzalez-Gil et al., 2018). Furthermore, He et al. have reported higher carbamazepine concentrations in the effluent than in the influent of WWTPs due to reactions transforming conjugates back to carbamazepine (He et al., 2019). Another plausible explanation of these results is that (2) removal percentage of an OMP is related to the amount of growth substrate added to the medium. Tran et al. reported that the removal percentage of diclofenac and carbamazepine was increased up to a maximum by adding higher concentrations of ammonium to activated sludge bottles (Tran et al., 2009). However, in our bioreactor, the concentration of growth substrates in the medium did not change, which results in a specific and limited number of enzymes and cofactors able to transform the selected pharmaceuticals. Oxygenases are key enzymes for the biodegradation of OMPs under aerobic conditions and require reducing power (i.e. NAD(P)H, FADH,) to transfer oxygen atoms to the respective substrate. When the substrate is used for growth, the cell first invests reducing power, but the subsequent catabolism of the transformation products restores the reducing power in the cell (Luo et al., 2014). In the case of co-metabolism, OMPs undergo only the first reaction with input of reducing power, which is energy costly for the cell (Cornelissen and Sijm, 1996). A previous study used a kinetic model to demonstrate that external donor and acceptor concentrations affect the removal percentage of OMPs because they change the NADH/NAD^+^ ratio inside the cell (Bae and Rittmann, 1995). Thus, a specific growth substrate concentration might be able to support only partial biotransformation of pharmaceuticals in WWTPs. On the other hand, Kennes-Veiga et al. did not observe a difference in OMP removal efficiency by increasing acetate concentrations in a continuous bioreactor (Kennes-Veiga et al., 2020). Instead, they only reported increasing biodegradation rate constants with increasing acetate loads. Furthermore, other studies observed different effects (increased, decreased, or unchanged) of growth substrate concentrations on biodegradation rate constants depending on the OMP (Liang et al., 2019; Su et al., 2015; Zhang et al., 2019). Consequently, the co-metabolic effect of growth substrates on pharmaceutical removal might be compound specific and might reach a maximum or a decay phase when there is competition for the same enzyme.

### 3.4. Bacterial community changes

Biomass samples were taken from the inoculum and from the bioreactor at different time points reflecting different pharmaceutical micropollutant influent concentrations (4 nM, 200 nM, and 800 nM) for bacterial 16S rRNA gene amplicon sequencing. Two time points with 4 nM were selected to account for the bacterial community changes inherent to the bioreactor maintenance and not to the shift in pharmaceutical concentration. Furthermore, technical triplicates were independently analysed for each time point.

Figure 3 shows the relative 16S rRNA gene abundance at the phylum level in the inoculum and bioreactor. First of all, the inoculum had a different bacterial community composition than the biomass in the bioreactor, where *Nitrospirae* and *Planctomycetes* increased in abundance and the rest decreased (i.e. *Actinobacteria* and *Chloroflexi*) or remained similar (i.e. *Proteobacteria* and *Acidobacteria*). Furthermore, the increase of pharmaceutical concentration in the bioreactor also induced a community shift. Overall, we saw an increase in *Acidobacteria* and *Bacteroidetes* and a decrease in *Nitrospirae* and *Planctomycetes* phyla. However, samples taken at two different time points (July and August) but with the same influent pharmaceutical concentration (4 nM) show a decrease in *Nitrospirae*, which suggests that this phylum is highly sensitive to small fluctuations during the bioreactor operation. On the other hand, a great decrease of around 50 % in *Nitrospirae* relative abundance at the highest pharmaceutical concentration points towards a putative toxic effect. Similarly, *Planctomycetes* also showed a slight shift in the two time points with the same concentrations, but ultimately, increasing pharmaceutical concentrations removed *Planctomycetes* to undetectable levels.

**Figure 3.**
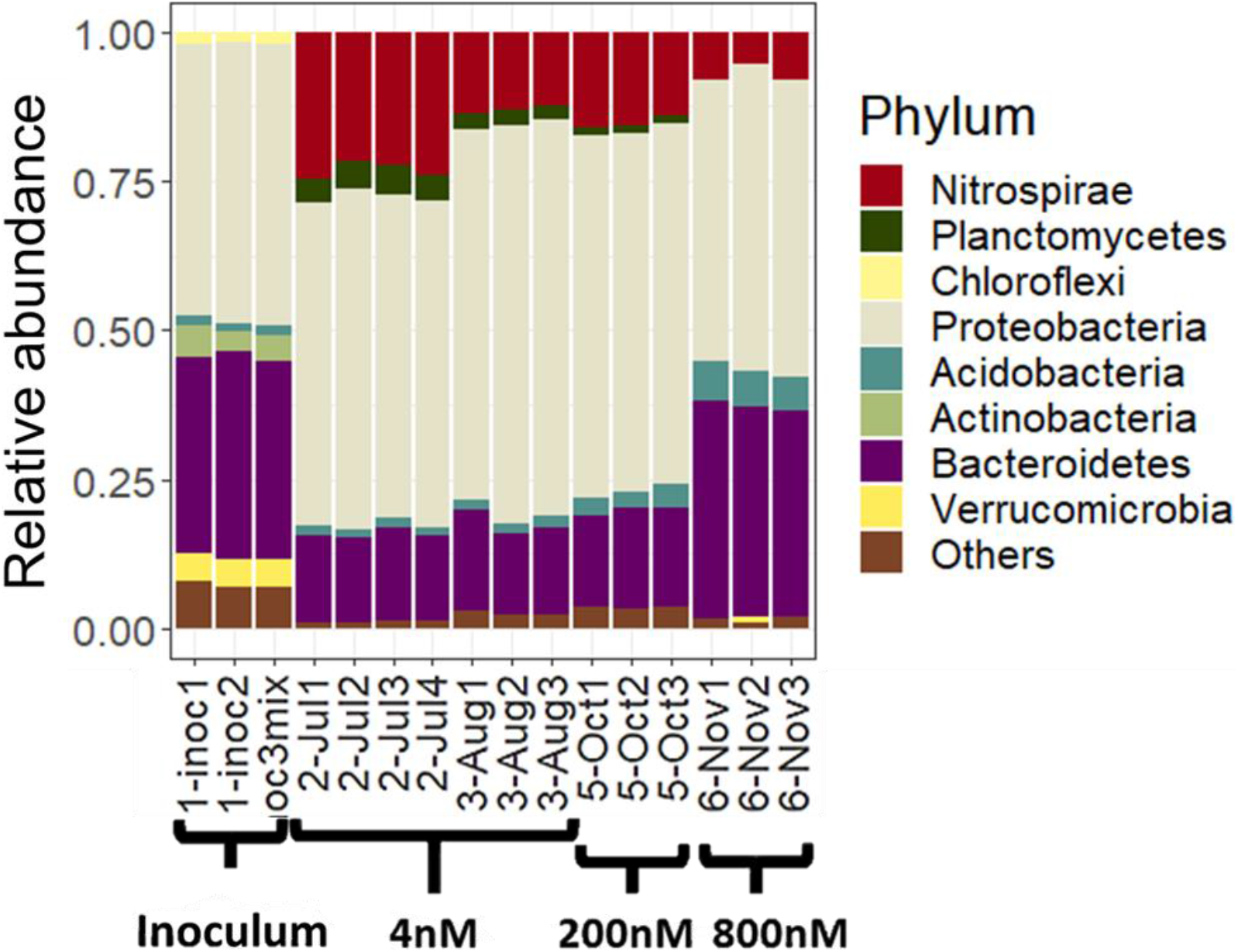
Relative abundance of 16S rRNA genes in the inoculum and bioreactor at different time points and different pharmaceutical concentrations added in the influent.

The relative abundance of ammonia oxidizing bacteria (AOB) significantly decreased with increasing pharmaceutical concentrations (data not shown). AOB consisted mainly of members of the *Nitrosomonadaceae* family and they considerably decreased in relative abundance in the bioreactor compared to the inoculum, in contrast to *Nitrospirae*. Due to the low abundance of AOB (less than 0.3%), the *Nitrospirae* might include complete ammonia oxidizing (comammox) species in addition to canonical nitrite-oxidizers (van Kessel et al., 2015). Since it is not possible to distinguish between comammox and nitrite-oxidizing *Nitrospira* using 16S rRNA gene sequencing methods (Koch et al., 2019), a PCR was performed using specific primers for comammox *amoA* (Pjevac et al., 2017). Primers targeting the clade A comammox *amoA* gene produced a single band of approximately 500 base pairs coinciding with the theoretical fragment length amplified by the primers. However, primers targeting clade B produced several bands not corresponding to the desired fragment. Therefore, we can conclude that clade A comammox may be present in the bioreactor and coexisted with AOB.

Previous studies reported a putative toxic effect of several pharmaceuticals towards *Nitrospira* and *Nitrosomonas* spp. (Bian et al., 2020; Dokianakis et al., 2004; Kraigher et al., 2008; Park and Seungdae, 2020; Wang and Gunsch, 2011a; b). *Nitrosomonas* spp. resisted higher concentrations of paracetamol and triclocarban than *Nitrospira* spp. (Bian et al., 2020; Park and Seungdae, 2020). Furthermore, heterotrophic activity for chemical oxygen demand removal was more robust than AOB activity in the presence of gemfibrozil and naproxen (Wang and Gunsch, 2011b). Wang et al. also reported that these two individual pharmaceuticals did not have an effect on ammonium removal, while the mixture of them inhibited nitrification and altered AOB community composition. In conclusion, each pharmaceutical has a different effect on *Nitrospira* and *Nitrosomonas* sp. and this effect is dose-dependent and stronger in a mixture. In our membrane bioreactor *Nitrospira* and *Nitrosomonas* spp. significantly decreased in relative abundance, but no effect was observed on nitrification. However, if there was inhibition it might not have been noticed due to the high HRT applied (3.5 days).

*Acidobacteria* and *Bacteroidetes* were the two phyla that increased in relative abundance with increasing pharmaceutical concentration. These phyla did not change in abundance at the two time points where the bioreactor had the same influent concentration of pharmaceuticals. Therefore, the relative abundance changes can be directly associated to the pharmaceutical concentration increase. Higher *Bacteroidetes* abundance has been previously associated to increased concentrations of pharmaceuticals and other OMPs in activated sludge reactors and river sediments (Drury et al., 2013; Jiang et al., 2017). Furthermore, there are several *Bacteroidetes* species able to degrade complex organic molecules (Kwon et al., 2019). Some studies have also associated *Acidobacteria* to micropollutant removal in soil and river sediments (George et al., 2009; Njeru et al., 2017; Nogales et al., 1999; Posselt et al., 2020). It is important to mention that an increase in abundance of these two phyla does not necessarily mean that they degrade pharmaceuticals. They could also be more resistant to toxic effects than other taxa giving them a growth advantage.

The genus *Dokdonella* (*Proteobacteria*) significantly increased in abundance from 0.5 % to 17 % of the total relative abundance in the microbial community at the highest pharmaceutical concentration in the bioreactor (800nM) (Figure S3). *Dokdonella* spp. consist of aerobic denitrifiers (Pishgar et al., 2019) and, in addition, they have been previously suggested to be responsible for the degradation of acetaminophen (Palma et al., 2018). In another study with activated sludge, *Dokdonella* spp. increased in abundance after adding around 400 μM of ibuprofen and 1 μM of tetracycline, so they might degrade other pharmaceuticals as well (Abdelrahman et al., 2018). Furthermore, this genus has been previously suggested as a pesticide and polycyclic hydrocarbon degrader (Bacosa and Inoue, 2015; Qi and Wei, 2017; Romero et al., 2019). Many other taxa (i.e. *Pseudomonas*, *Bacillus*, *Flavobacterium*, *Ensifer*, *Delftia* spp.) have been isolated from activated sludge and soil and have been shown to degrade acetaminophen in pure and mixed cultures (Akay and Tezel, 2020; Chopra and Kumar, 2020; Park and Oh, 2020; Żur et al., 2018). In our bioreactor, some of these genera were not present and others did not show an increase with higher pharmaceutical concentrations (data not shown). For this reason, we propose *Dokdonella* as the main acetaminophen degrader in our bioreactor experiment.

Alpha diversity was analysed with three different indices: Chao1, Shannon, and Simpson (Figure S4). The richness (number of different species), calculated with the Chao1 index, decreased in the bioreactor compared to the inoculum and it further decreased with the highest pharmaceutical concentration. The main driver of this change could be the synthetic medium used to feed the bioreactor: we simplified the influent organic matter to acetate and methanol compared to the large variety of carbon sources usually found in the influent of WWTPs. Furthermore, a decrease of richness with pharmaceutical concentration could be explained by a toxicity effect. Shannon index measures the richness and evenness (distribution of individuals among the species increases) of a sample whereas Simpson’s index of diversity (1 – Simpson index) gives more weight to the abundance and evenness of species (Lemos et al., 2011). Both indices considerably decreased in the bioreactor biomass compared to the inoculum. In the middle of the experiment, they increased again, and decreased during the highest pharmaceutical concentration. The samples taken at two consecutive time points at the same pharmaceutical concentration (4 nM) have very different evenness values, so this diversity characteristic cannot be associated to the pharmaceuticals. As seen in Figure S2, there are small fluctuations in the influent growth substrate concentrations that might be responsible of these changes. Previous studies have correlated biodiversity (richness and evenness) to higher biotransformation rates of specific micropollutants (Johnson et al., 2015; Stadler et al., 2018; Torresi et al., 2016). However, those studies did not correlate the diversity changes to the addition of micropollutants, which might increase diversity due to a putative cooperation of different taxa in order to degrade the chemicals (Vasiliadou et al., 2018) or decrease diversity due to putative toxic effects (Chonova et al., 2016).

### 3.5. Mobile genetic elements relative abundance

In this study, we tested whether increasing pharmaceutical concentrations in the bioreactor led to an increase in the relative abundance of IncP-1 plasmids and IS*1071* insertion sequences as a measure for MGEs. We performed a qPCR of the *trfA* replication gene of different IncP-1 plasmids and the *tnpA* transposase gene of the IS*1071* insertion sequence. As internal standard, the bacterial 16S rRNA gene was used. Interestingly, higher pharmaceutical concentrations did not significantly increase relative abundance of MGEs Figure S5). The studied MGEs remained stable in the bacterial community independent of the pharmaceutical concentration. This result does not necessarily mean that plasmids are not being transferred, as they can also remain in a dynamic equilibrium in the bioreactor. Yang et al. observed that the relative abundance of an external plasmid remained unchanged despite being transferred in a membrane reactor (Yang et al., 2013). The biomass density in the bioreactor presented in this study was ten times lower than a common WWTP activated sludge reactor. As conjugation events are more likely to happen at higher densities or in environments where cells are in close contact such as granules and biofilms (Rios-Miguel et al., 2020), this experiment might underestimate the true role that MGEs play in the degradation of pharmaceutical micropollutants. Furthermore, previous studies reported higher relative abundances of MGEs in sites with higher OMP exposure over much longer periods of time (years) compared to our bioreactor experiment that lasted 2.5 months (Dealtry et al., 2014; Dunon et al., 2013). These might be reasons why MGE relative abundances were stable in our bioreactor.

## 4. Conclusions

This study contributes to a better understanding of pharmaceutical removal limitations in activated sludge processes. The observed relative abundance shifts of the microbial community suggest toxic effects on specific phyla relevant to WWTP functioning and point out several microbial groups potentially involved in degrading pharmaceuticals. The main conclusions from our experimental data are:

1. Selected pharmaceuticals (diclofenac, metoprolol, metformin, carbamazepine, and fluoxetine) were not fully degraded under activated sludge conditions in a MBR. Only acetaminophen was fully removed.
2. Higher pharmaceutical influent concentration proportionally increased the removal rate of each compound.
3. The removal percentage was not correlated to influent concentration and remained relatively stable in the order: acetaminophen (100%) > fluoxetine (50%) > metoprolol (25%) > diclofenac (20%) > metformin (15%) > carbamazepine (10%).
4. Metformin removal increased to 80-100% when HRT or biomass concentration increased in the bioreactor. Removal % of the other pharmaceuticals could not be improved by extending the HRT nor increasing biomass concentration.
5. The microbial community changed in response to higher pharmaceutical concentrations: *Nitrospira* and *Planctomycetes* decreased and *Bacteroidetes* and *Acidobacteria* increased in relative abundance.
6. *Dokdonella* spp. could be the main acetaminophen degraders under activated sludge conditions.
7. Increasing pharmaceutical concentrations did not increase relative abundance of mobile genetic elements (IncP-1 plasmids and *IS1071*). The biomass density or the pharmaceutical concentration in the bioreactor might have been too low to increase conjugation.

## 5. Data availability statement

All sequencing data were submitted to the GenBank database under the BioProject ID PRJNA670630 (reviewer link: https://dataview.ncbi.nlm.nih.gov/object/PRJNA670630?reviewer=dt2eeti6d8l048a7r6t9efict4).The data that support the findings of this manuscript are available in Dans Easy with the following digital object identifier: 10.17026/dans-zhx-nuux (it will be accessible upon acceptance of the manuscript).

## 6. Authorship contribution statement

All authors started the project and contributed to the conceptual framework. ARM conducted the experiments and data analyses. ARM wrote the manuscript and all authors contributed to improve versions of the manuscript.

## 8. Acknowledgements

The authors thank the CER-CEC group (Tamara van Bergen, Jan Hendriks, Tom Nolte, Rosalie van Zelm, Ad Ragas) for discussion, Guylaine Nuijten for the technical support with bioreactors, the Radboud UMC team (Martien Graumans, Rob Anzion, Maurice van Dael, and Paul Scheepers) for help with the LC-MS measurements, Rob de Graaf for help with acetate measurements, Pieter Blom for help with N-measurements and comammox PCR, Hanna Koch for discussion about *Nitrospira*, Carmen Hogendoorn for help with methanol measurements, Stefanie Berger for help with molecular analyses, and Lianna Pogoshyan and Anniek de Jong for help with DADA2. This work was supported by the NWO-domain TTW [grant number 15759], the Netherlands, and the SIAM Gravitation grant funded by NWO [grant number 024.002.002].

