## Supplementary information for "Effect of concentration and hydraulic reaction time on the removal of pharmaceutical compounds in a membrane bioreactor inoculated with activated sludge"

### Table S1. Bioreactor medium composition

| **Compound** | **Concentration** |
| --- | --- |
| **Bioreactor medium** | |
| NH_4_Cl | 3.57 mM |
| NaAcetate x 3 H_2_O | 3.14 mM |
| KH_2_PO_4_ | 0.16 mM |
| MgSO_4_ x 7H_2_O | 0.04 mM |
| CaCl_2_ | 0.09 mM |
| NaHCO_3_ | 5.95 mM |
| TES I | 1 mL/L |
| TES II | 1 mL/L |
| Pharmaceutical solution in methanol/ demi water (1:1) | 1 mL/L (from stocks with different concentrations: 40 µM to 8 mM) |
| **Trace Elements Solution I (1000x)** | |
| NTA (Trisodium Nitrilotriacetate), pH 8 (NaOH) | 38.9 mM |
| FeSO_4_ x 7 H_2_O | 17.39 mM |
| **Trace Elements Solution II (1000x)** | |
| NTA (Trisodium Nitrilotriacetate), pH 8 (NaOH) | 58.35 mM |
| ZnSO_4_ x 7 H_2_O | 1.5 mM |
| CoCl_2_ x 6 H_2_O | 1.01 mM |
| MnCl_2_ x 4 H_2_O | 5 mM |
| CuSO_4_ x 5 H_2_O | 1 mM |
| NaMoO_4_ x 2 H_2_O | 0.91 mM |
| NiCl_2_ x 6 H_2_O | 0.8 mM |
| NaSeO_4_ x 10 H_2_O | 0.61 mM |
| H_3_BO_4_ | 1.8 mM |
| CeCl_3_ x 6 H_2_O | 0.68 mM |

### Table S2. List of primers used in this study

| Name primer | Sequence | Target gene | Amplified fragment size | Reference |
| --- | --- | --- | --- | --- |
| Bac341F | 5’-CCTACGGGNGGCWGCAG-3’ | 16S rRNA gene for all bacteria | 464 bp | (Klindworth et al., 2013) |
| Bac785R | 5’-GACTACHVGGGTATCTAATCC-3’ |  |  |  |
| ComaA244F | 5’-TAYAAYTGGGTSAAYTA-3’ | *amoA* gene for clade A comammox | 415 bp | (Pjevac et al., 2017) |
| ComaA659R | 5’-ARATCATSGTGCTRTG-3’ |  |  |  |
| ComaB244F | 5’-TAYTTCTGGACRTTYTA-3’ | *amoA* gene for clade B comammox |  |  |
| ComaB659R | 5’-ARATCCARACDGTGTG-3’ |  |  |  |
| trfA_αβε_F | 5’-TTCACSTTCTACGAGMTKTGCCAGGAC-3’ | trfA gene of IncP-1 α, β, and ε plasmids | 281 bp | (Bahl et al., 2009) |
| trfA_αβε_R | 5’-GWCAGCTTGCGGTACTTCTCCCA-3’ |  |  |  |
| trfA_γ_F | 5’-TTCACTTTTTACGAGCTTTGCAGCGAC-3’ | trfA gene of IncP-1 γ plasmids |  |  |
| trfA_γ_R | 5’-GTCAGCTCGCGGTACTTCTCCCA-3’ |  |  |  |
| trfA_δ_F | 5’-TTCACGTTCTACGAGCTTTGCACAGAC-3’ | trfA gene of IncP-1 δ plasmids |  |  |
| trfA_δ_R | 5’-GACAGCTCGCGGTACTTTTCCCA-3’ |  |  |  |
| tnpA_F | 5’-GCTTGGTCACTTCTGGGTCTTC-3’ | tnpA gene of IS*1071* | 180 bp | (Providenti et al., 2006) |
| tnpA_R | 5’-CTATGCCCGTCTATCGTTACCC |  |  |  |

### Table S3. Bioreactor monitoring

| **Measurement** | **Figure** |
| --- | --- |
| **Total suspended solids** | Figure S1 |
| **Ammonium, nitrite, nitrate, acetate, and methanol** | Figure S2 |
| **Pharmaceuticals** | Figures 1,2 |
| **16S rRNA gene** | Figures 3, S3,S4 |
| **Mobile genetic elements** | Figure S5 |

### Figure S1. Total suspended solids values along the bioreactor acclimatization period and experiments.

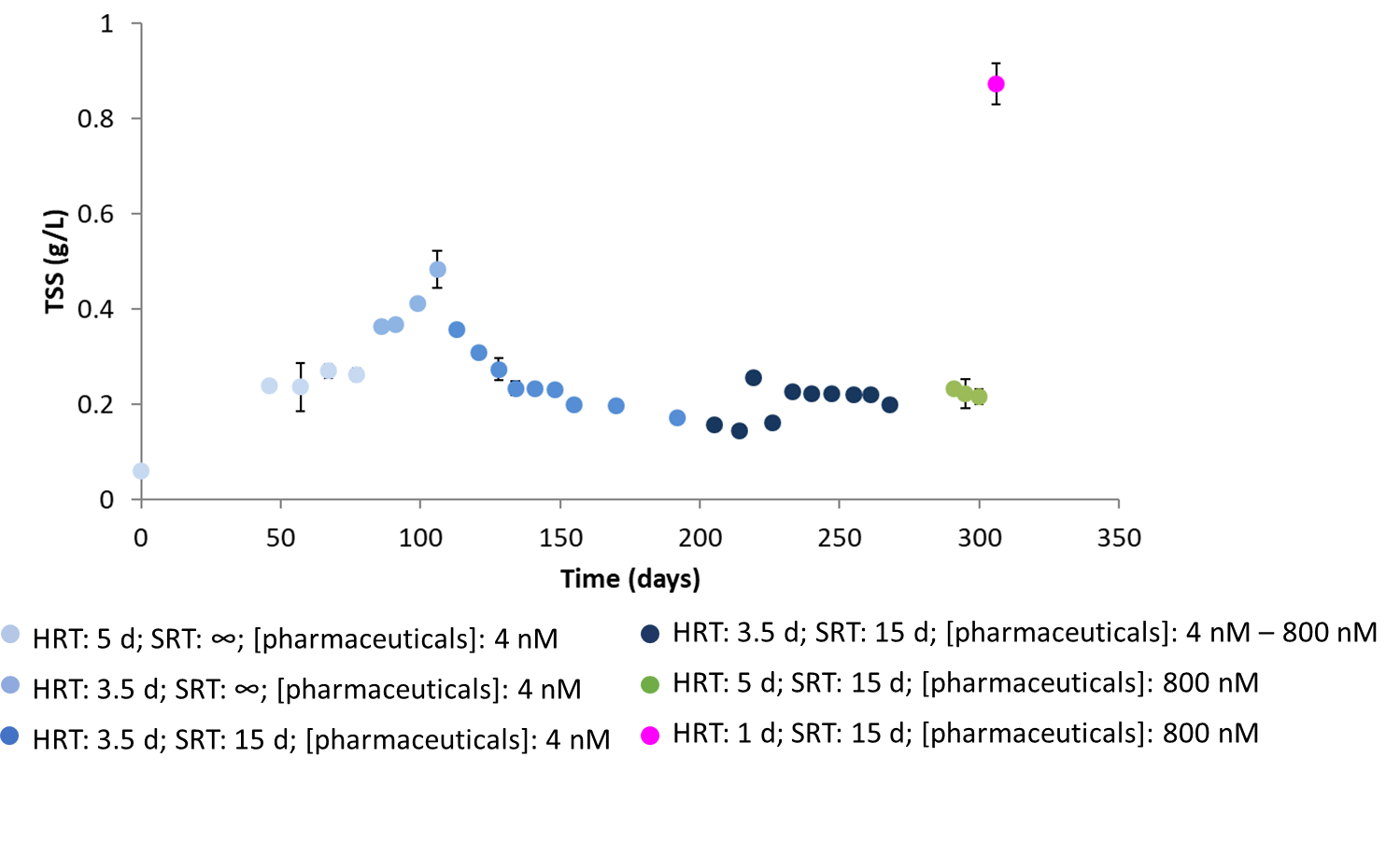

### Figure S2. Carbon and nitrogen measurements in the influent (A) and effluent (B) along the bioreactor experiment

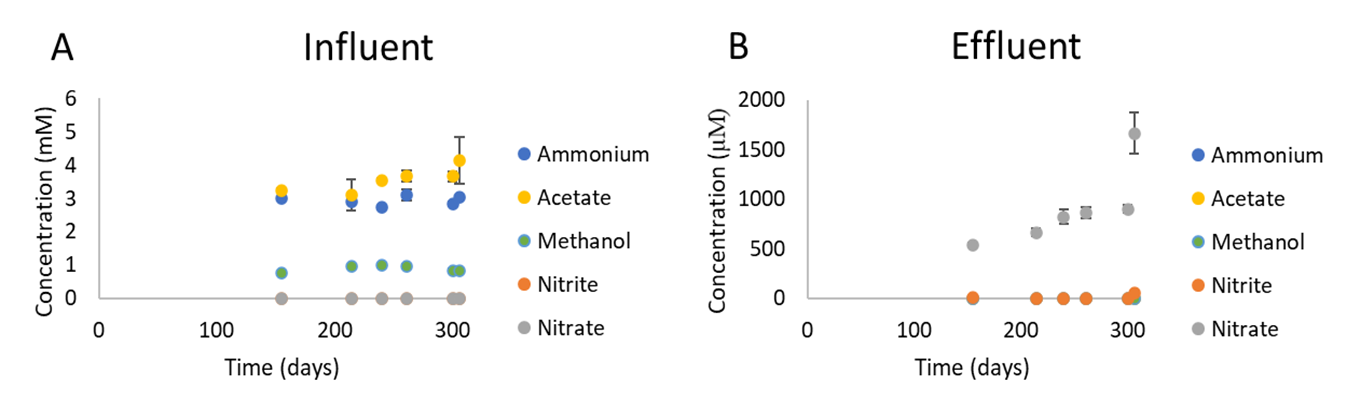

Figure S2. Carbon and nitrogen measurements during the bioreactor experiment. The conditions in the bioreactor at the six time points shown in the graph were: Day 155: HRT = 3.5 days, [Pharmaceuticals] = 4 nM; Day 214: HRT = 3.5 days, [Pharmaceuticals] = 40 nM; Day 240: HRT = 3.5 days, [Pharmaceuticals] = 200 nM; Day 261: HRT = 3.5 days, [Pharmaceuticals] = 800 nM; Day 300: HRT = 5 days, [Pharmaceuticals] = 800 nM; Day 306: HRT = 1 day, [Pharmaceuticals] = 800 nM.

### Figure S3. Relative abundance of the *Dokdonella* genus in the bioreactor at different time points and pharmaceutical concentrations

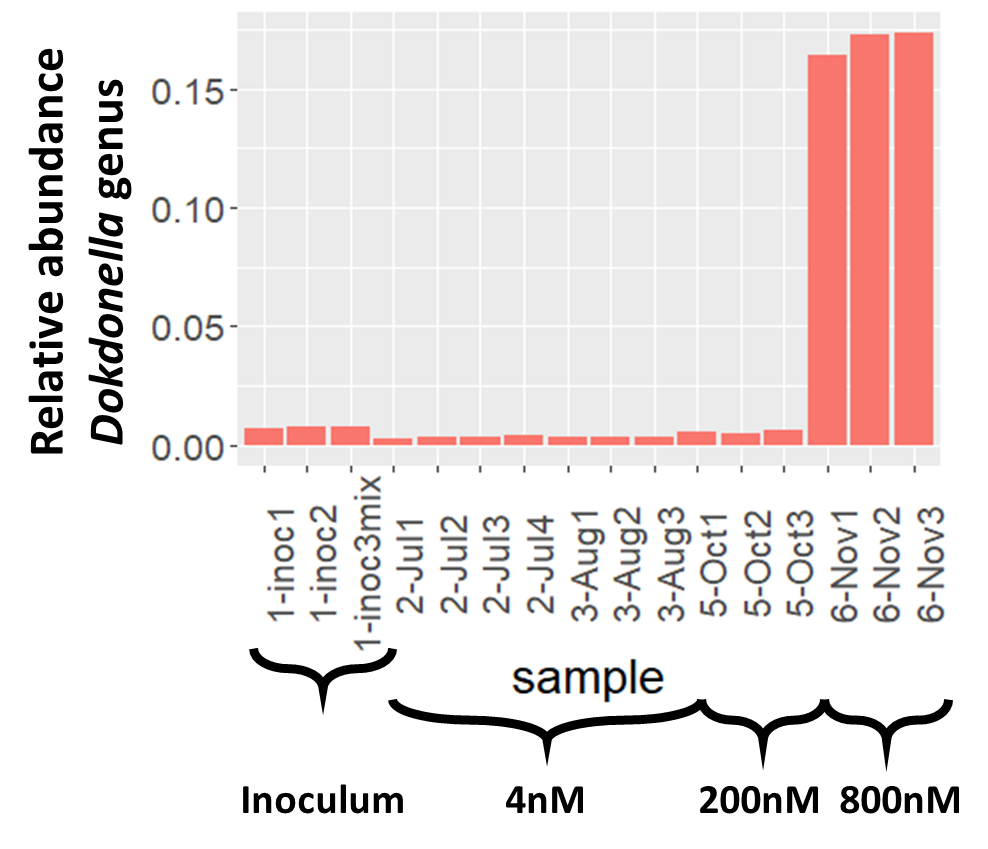

### Figure S4. Alpha diversity in the inoculum and bioreactor at different time points

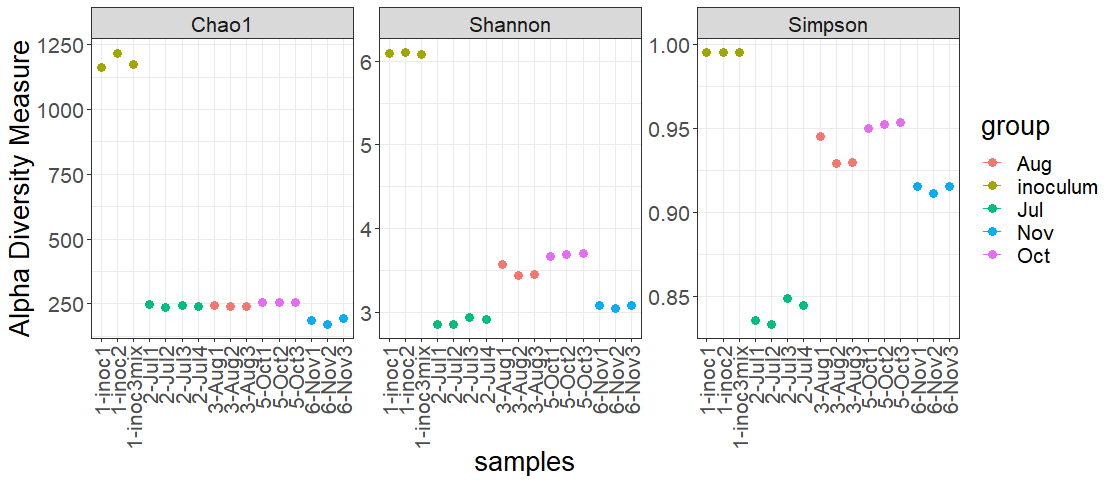

### Figure S5. Mobile genetic elements genes *versus* bacterial 16S rRNA gene relative abundance in the bioreactor when pharmaceutical concentrations were 4 nM and 800 nM.

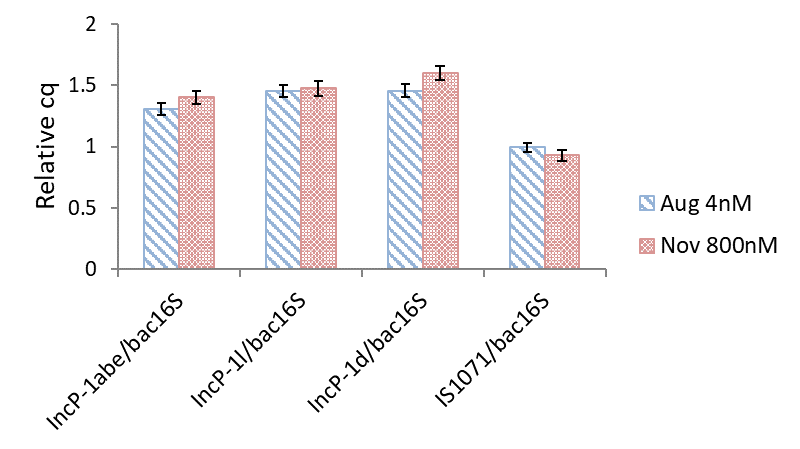
